# Endothelial Pannexin 1–TRPV4 channel signaling lowers pulmonary arterial pressure

**DOI:** 10.1101/2021.03.09.434532

**Authors:** Zdravka Daneva, Matteo Ottolini, Yen-Lin Chen, Eliska Klimentova, Soham A. Shah, Richard D. Minshall, Cheikh I. Seye, Victor E. Laubach, Brant E. Isakson, Swapnil K. Sonkusare

## Abstract

Pannexin 1 (Panx1) is an ATP-efflux channel that controls endothelial function in the systemic circulation. However, the roles of endothelial Panx1 in resistance-sized pulmonary arteries (PAs) are unknown. Extracellular ATP dilates PAs through activation of endothelial TRPV4 (transient receptor potential vanilloid 4) ion channels. We hypothesized that endothelial Panx1–ATP– TRPV4 channel signaling promotes vasodilation and lowers pulmonary arterial pressure (PAP). Endothelial, but not smooth muscle, knockout of Panx1 or TRPV4 increased PA contractility and raised PAP. Panx1-effluxed extracellular ATP signaled through purinergic P2Y2 receptor (P2Y2R) to activate protein kinase Cα (PKCα), which in turn activated endothelial TRPV4 channels. Finally, caveolin-1 provided a signaling scaffold for endothelial Panx1, P2Y2R, PKCα, and TRPV4 channels in PAs, promoting their spatial proximity and enabling signaling interactions. These results indicate that endothelial Panx1–P2Y2R–TRPV4 channel signaling, facilitated by caveolin-1, reduces PA contractility and lowers PAP.

## Introduction

The pulmonary endothelium exerts a dilatory influence on small, resistance-sized pulmonary arteries (PAs) and thereby lowers pulmonary arterial pressure (PAP). However, endothelial signaling mechanisms that control PA contractility remain poorly understood. In this regard, pannexin 1 (Panx1), which is expressed in the pulmonary endothelium and epithelium^1^, has emerged as a crucial controller of endothelial function^2, 3^. Panx1, the most studied member of the pannexin family, forms a hexameric transmembrane channel at the cell membrane that allows efflux of ATP from the cytosol^4, 5^. Previous studies have indicated that Panx1_EC_ promotes endothelium-dependent dilation of systemic arteries^6, 7^, and endothelial cell (EC) Panx1 (Panx1_EC_) has been linked to inflammation in pulmonary capillaries^8^. Beyond this, however, the physiological roles of Panx1_EC_ in the pulmonary vasculature are largely unknown.

Extracellular ATP (eATP) was recently shown to activate TRPV4 (transient receptor potential vanilloid 4) channels in the endothelium of small PAs^9^, establishing endothelial TRPV4 (TRPV4_EC_) channels as potential signaling targets of Panx1_EC_ in the pulmonary circulation. Ca^2+^ influx through TRPV4_EC_ channels is known to dilate small PAs through activation of endothelial nitric oxide synthase (eNOS)^9^. These observations suggest that Panx1_EC_-released eATP may act through TRPV4_EC_ channels to reduce PA contractility and lower PAP.

Purinergic receptor signaling is an essential regulator of pulmonary vascular function^10–13^. Previous studies in small PAs showed that eATP activates TRPV4_EC_ channels through P2 purinergic receptors, although the precise P2 receptor subtype was not identified^9^. Pulmonary endothelium expresses both P2Y and P2X receptor subtypes. Konduri et al. showed that eATP dilates PAs through P2Y2 receptor (P2Y2R) activation and subsequent endothelial NO release^13^. Recent evidence from systemic ECs and other cell types also supports P2Y2R-dependent activation of TRPV4 channels by eATP^14, 15^. These findings raise the possibility that the endothelial P2Y2 receptor (P2Y2R_EC_) may be the signaling intermediate for Panx1_EC_–TRPV4_EC_ channel communication in PAs.

The linkage between Panx1_EC_-mediated eATP release and subsequent activation of P2Y2R_EC_–TRPV4_EC_ signaling could depend on the spatial proximity of individual elements— Panx1_EC_, P2Y2R_EC_, and TRPV4_EC_—a functionality possibly provided by a signaling scaffold. Caveolin-1 (Cav-1), a structural protein that interacts with and stabilizes other proteins in the pulmonary circulation^16^, co-localizes with Panx1, P2Y2R, and TRPV4 channels in multiple cell types^17–19^. Notably, global Cav-1^-/-^ mice show elevated PAP, and endothelial Cav-1 (Cav-1_EC_)-dependent signaling is impaired in pulmonary hypertension^20–22^.

Here, we tested the hypothesis that Panx1_EC_–P2Y2R_EC_–TRPV4_EC_ channel signaling, supported by a signaling scaffold provided by Cav-1_EC_, reduces PA contractility and PAP. Using inducible, EC-specific Panx1^-/-^, TRPV4^-/-^, P2Y2R^-/-^ and Cav-1_EC_^-/-^ mice, we show that endothelial Panx1–P2Y2R–TRPV4 signaling reduces PA contractility and lowers PAP. Panx1_EC_-generated eATP acts via P2Y2R_EC_ stimulation to activate protein kinase Cα (PKCα) and thereby increase TRPV4_EC_ channel activity. Panx1_EC_, P2Y2R_EC_, PKCα, and TRPV4_EC_ channels co-localize with Cav-1_EC,_ ensuring spatial proximity among the individual elements and supporting signaling interactions. Overall, these findings advance our understanding of endothelial mechanisms that control PAP and suggest the possibility of targeting these mechanisms to lower PAP in pulmonary vascular disorders.

## Results

### Endothelial, but not smooth muscle, Panx1**–**TRPV4 signaling lowers PA contractility

To clearly define the physiological roles of Panx1_EC_ and TRPV4_EC_ channels, we utilized tamoxifen-inducible, EC-specific Panx1_EC_^-/-^ and TRPV4_EC_^-/-^ mice^23, 24^. Tamoxifen-injected TRPV4^fl/fl^ Cre^-^ (TRPV4^fl/fl^) or Panx1^fl/fl^ Cre^-^ (Panx1^fl/fl^) mice were used as controls^8, 23^. TRPV4_EC_^-/-^ mice showed elevated right ventricular systolic pressure (RVSP), a commonly used *in vivo* indicator of PAP (Fig. 1A). In pressure myography experiments, ATP (1 μmol/L)-induced dilation was absent in PAs from TRPV4_EC_^-/-^ mice (Fig. 1B), confirming that ATP dilates PAs through TRPV4_EC_ channels. RVSP was also elevated in Panx1_EC_^-/-^ mice (Fig. 1C). The Fulton Index, a ratio of right ventricular (RV) weight to left ventricle plus septal (LV + S) weight, was not altered in TRPV4_EC_^-/-^ or Panx1_EC_^-/-^ mice compared with the respective control mice, suggesting a lack of right ventricular hypertrophy in these mice (Table 1). Importantly, baseline RVSP was not altered in inducible, SMC-specific TRPV4 (TRPV4_SMC_^-/-^) or Panx1 (Panx1_SMC_^-/-^) knockout mice (Fig. 1A and C). Functional cardiac MRI studies indicated no alterations in cardiac function in TRPV4_EC_^-/-^ or Panx1_EC_^-/-^ mice compared with the respective control mice (Table 1), suggesting that the changes in RVSP were not due to altered cardiac function.

**Figure 1.**
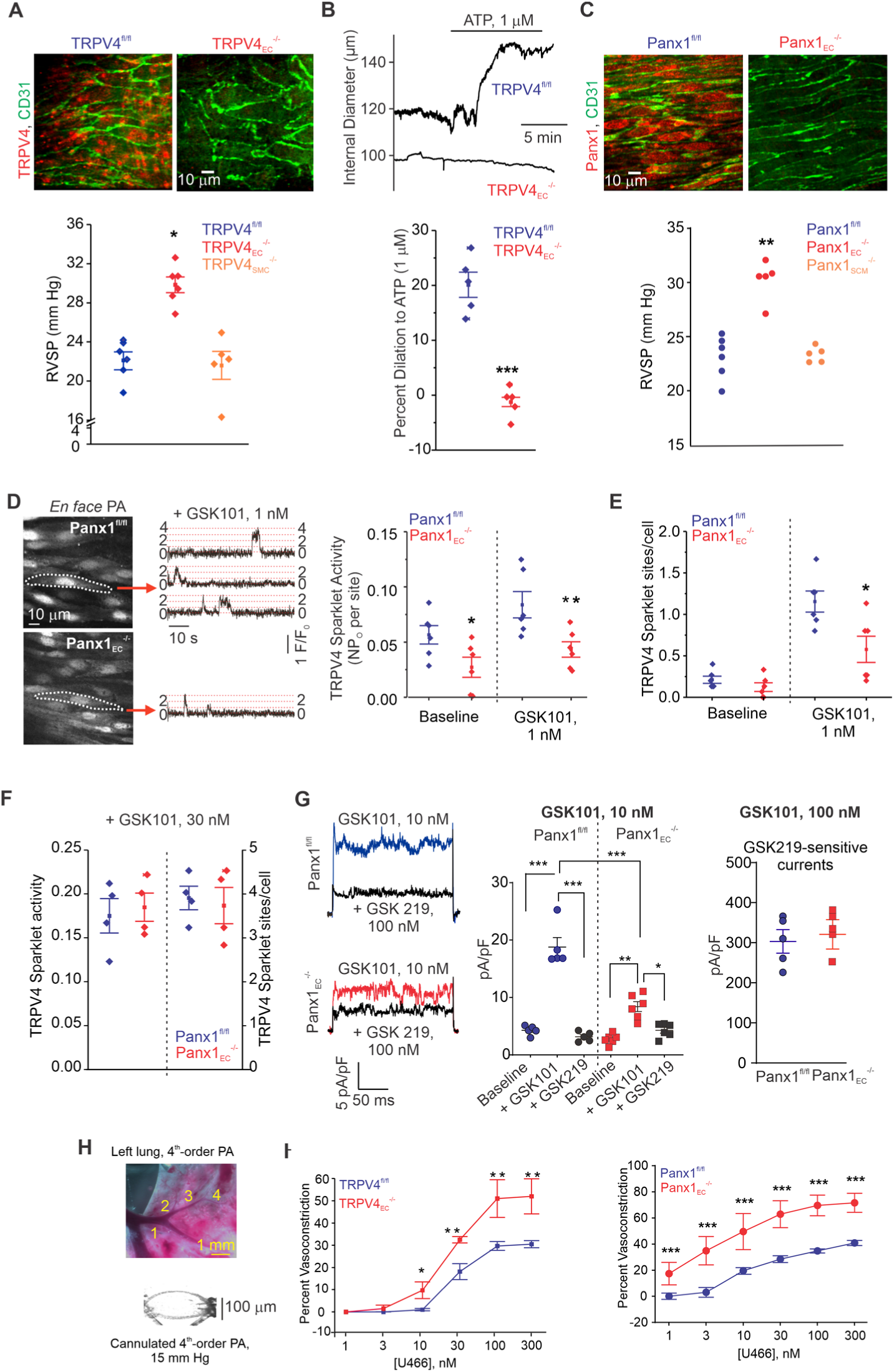
Panx1_EC_–TRPV4_EC_ signaling reduces PA contractility and lowers PAP. **A**, *top,* Immunofluorescence images of *en face* 4^th^-order PAs from TRPV4^fl/fl^ (*left*) and TRPV4_EC_^-/-^ (*right*) mice. CD31 immunofluorescence indicates endothelial cells. *Bottom,* Average resting RVSP values in TRPV4^fl/fl^, TRPV4_EC_^-/-^, and TRPV4_SMC_^-/-^ mice (n = 6; \**P <* 0.05 vs. TRPV4^fl/fl^; one-way ANOVA). **B,** *top,* Representative diameter traces showing ATP (1 μmol/L)-induced dilation of PAs from TRPV4^fl/fl^ and TRPV4_EC_^-/-^ mice, pre-constricted with the thromboxane A2 receptor analog U46619 (50 nmol/L). Fourth-order PAs were pressurized to 15 mm Hg. *Bottom,* Percent dilation of PAs from TRPV4^fl/fl^ and TRPV4_EC_^-/-^ mice in response to ATP (1 μmol/L; n = 5–10; \*\*\**P <* 0.01 vs. TRPV4^fl/fl^ [ATP 1 μmol/L]; t-test). **C,** *top,* Immunofluorescence images of *en face* 4^th^-order PAs from Panx1^fl/fl^ (*left*) and Panx1_EC_^-/-^ (*right*) mice. *Bottom,* Average resting RVSP values in Panx1^fl/fl^ Panx1 and Panx1_SMC_ mice (n = 5; \*\**P <* 0.01 vs. Panx1; one-way ANOVA). **D,** *left,* Grayscale image of a field of view of an *en face* preparation of Fluo-4–loaded PAs from Panx1^fl/fl^ and Panx1_EC_^-/-^ mice showing approximately 20 ECs. Dotted areas indicate TRPV4_EC_ sparklet sites (20 μmol/L CPA + 10 nmol/L GSK101). *Center,* Representative traces showing TRPV4_EC_ sparklet activity in *en face* preparations of PAs from Panx1^fl/fl^ and Panx1_EC_^-/-^ mice in response to GSK101 (10 nmol/L). Experiments were performed in Fluo-4–loaded PAs in the presence of CPA (20 μmol/L), included to eliminate Ca^2+^ release from intracellular stores. *Right,* TRPV4_EC_ sparklet activity (NP_O_) per site in *en face* preparations of PAs from Panx1^fl/fl^ and Panx1_EC_^-/-^ mice under baseline conditions (i.e., 20 μmol/L CPA) and in response to 1 nmol/L GSK101 (n = 6; \**P <* 0.05, \*\**P <* 0.01 vs. Panx1^fl/fl^; two-way ANOVA). ‘N’ is the number of channels per site and ‘P_O_’ is the open state probability of the channel. **E,** TRPV4_EC_ sparklet activity, expressed as sites per cell, in *en face* preparations of PAs from Panx1^fl/fl^ and Panx1_EC_^-/-^ mice under baseline conditions (i.e., 20 μmol/L CPA) and in response to 1 nmol/L GSK101 (n = 6; \**P <* 0.05 vs. Panx1^fl/fl^; two-way ANOVA). **F,** TRPV4_EC_ sparklet activity (NP_O_) per site and TRPV4 sparklet sites per cell in *en face* preparations of PAs from Panx1^fl/fl^ and Panx1_EC_^-/-^ mice in response to 30 nmol/L GSK101 (n = 6). **G,** *left,* representative GSK101 (10 nmol/L)–induced outward TRPV4_EC_ currents in freshly isolated ECs from Panx1^fl/fl^ or Panx1_EC_^-/-^ mice and effect of GSK2193874 (GSK219, TRPV4 inhibitor, 100 nmol/L) in the presence of GSK101, currents were elicited by a 200 ms voltage step from −50 mV to +100 mV; *center,* scatterplot showing outward currents at +100 mV under baseline conditions, after the addition of GSK101 (10 nM), and after the addition of GSK219 (100 nM), n=5-6 cell, one-way ANOVA, *right*, scatterplot showing GSK219-sensitive TRPV4_EC_ currents in the presence of GSK101 (100 nmol/L; n = 5). **H,** *top,* an image showing the left lung and the order system used to isolate 4^th^ order PAs in this study; *bottom,* an image of a 4^th^ order PA cannulated and pressurized at 15 mm Hg. **I,** *left,* Percent constriction of PAs from TRPV4^fl/fl^ and TRPV4_EC_^-/-^ mice in response to U46619 (1–300 nmol/L) (n = 5; \**P* < 0.05, \*\**P <* 0.01 vs. Panx1^fl/fl^; two-way ANOVA). *Right,* Percent constriction of PAs from Panx1^fl/fl^ and Panx1_EC_^-/-^ mice in response to U46619 (1–300 nmol/L; n = 5; \*\*\**P <* 0.001 vs. Panx1^fl/fl^; two-way ANOVA).

**Table 1.**
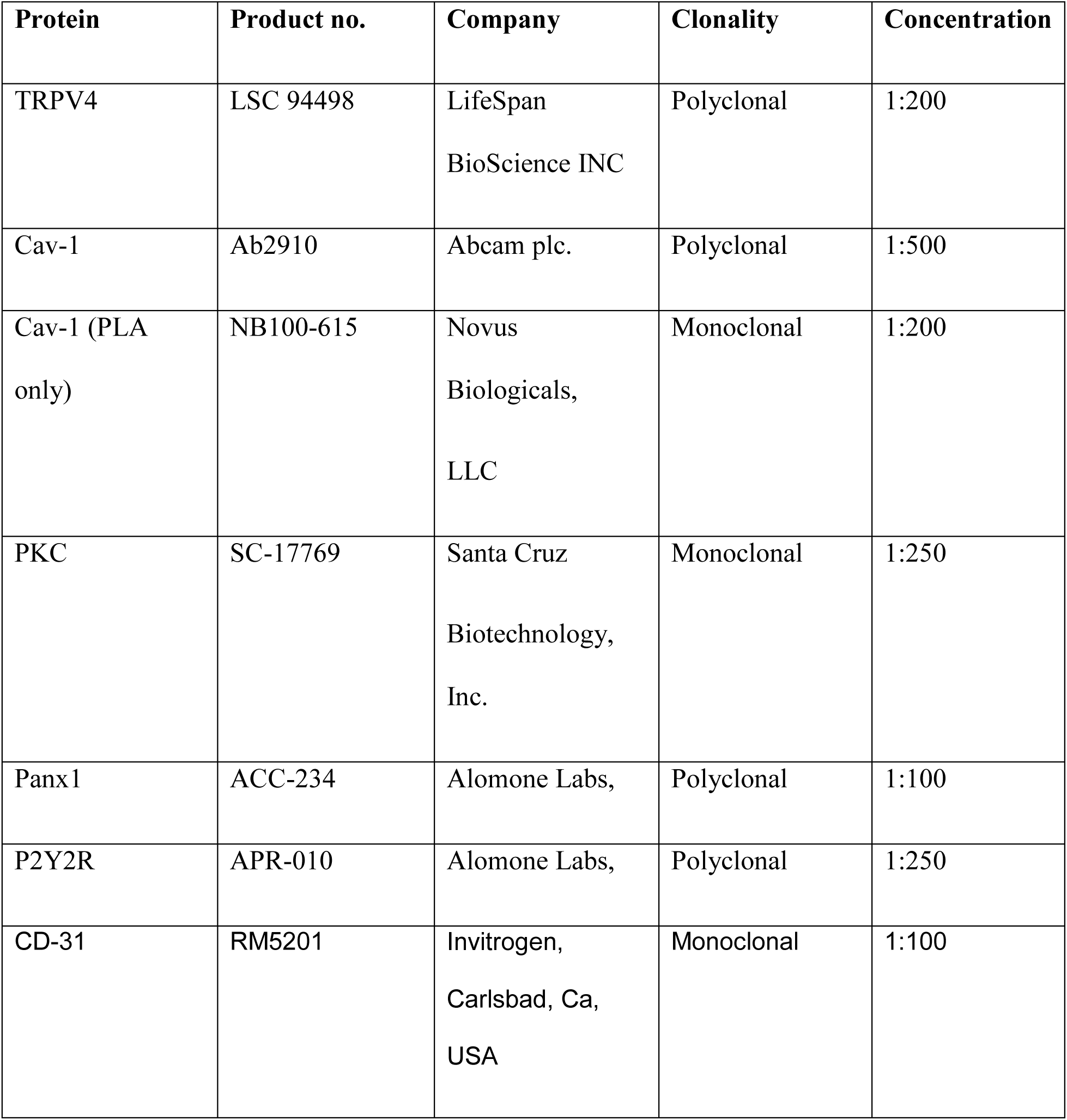
Fulton Index and Functional MRI analysis of cardiac function in TRPV4^fl/fl^, TRPV4_EC_^-/-^, Panx1^fl/fl^ and Panx1_EC_^-/-^ mice. Average Fulton Index, end diastolic and systolic volume (EDV and ESV; μL), ejection fraction (EF; %), stroke volume (SV; μL), R-R interval (ms), and cardiac output (CO; mL/min). Data are presented as means ± SEM (n = 5–8 mice).

Localized, unitary Ca^2+^ influx signals through TRPV4_EC_ channels, termed *TRPV4_EC_ sparklets*^25^, were recorded in *en face*, 4th-order PAs (∼ 50 μm) loaded with Fluo-4. Baseline TRPV4_EC_ sparklet activity and activity induced by a low concentration (1 nmol/L) of the specific TRPV4 channel agonist, GSK1016790A (hereafter, GSK101), were significantly reduced in PAs from Panx1_EC_^-/-^ mice compared with those from Panx1^fl/fl^ mice (Fig. 1D). Additionally, the number of TRPV4_EC_ sparklet sites per cell was decreased in PAs from Panx1_EC_^-/-^ mice (Fig. 1E). At a higher level of TRPV4 channel activation (30 nmol/L GSK101), sparklet activity per site and sparklet sites per cell were not different between Panx1_EC_^-/-^ and control (Panx1^fl/fl^) mice (Fig. 1F). Further, outwards currents through TRPV4_EC_ channels, elicited by 10 nM GSK101, were also lower in Panx1_EC_^-/-^ than Panx1^fl/fl^ mice (Fig. 1G, *left* and *center*). However, TRPV4_EC_ channel currents, elicited by 100 nM GSK101, were not different between Panx1_EC_^-/-^ and Panx1^fl/fl^ mice (Fig. 1G, *right*). These data support the concept that the reduced TRPV4_EC_ channel activity in Panx1_EC_^-/-^ mice is due to impaired channel regulation rather than a decrease in the number of functional TRPV4_EC_ channels. Moreover, isolated, pressurized 4^th^-order PAs (Fig. 1H) from TRPV4_EC_^-/-^ mice and Panx1_EC_^-/-^ mice exhibited a greater contractile response to the thromboxane A_2_ receptor agonist U46619 (1–300 nmol/L; Fig. 1I). Together, these data provide the first evidence that Panx1_EC_, via regulation of TRPV4_EC_ channel activity, lowers resting PAP.

### Panx1_EC_-generated eATP acts through purinergic P2Y2R_EC_ stimulation to activate TRPV4_EC_ channels

Bioluminescence measurements confirmed lower baseline eATP levels in PAs from Panx1_EC_^-/-^ mice compared with PAs from Panx1^fl/fl^ mice (Fig. 2A), supporting an essential role for Panx1_EC_ channels as an eATP-release mechanism in PAs. PAs from TRPV4_EC_^-/-^ mice, however, exhibited unaltered basal eATP levels. eATP was recently identified as a novel endogenous activator of TRPV4_EC_ channels in the pulmonary circulation^9^. Therefore, we tested whether Panx1_EC_ activates TRPV4_EC_ channels via eATP release. Addition of the eATP-hydrolyzing enzyme apyrase (100 U/mL) reduced the activity of TRPV4_EC_ sparklets in PAs from control mice but not those from Panx1_EC_^-/-^ mice (Fig. 2B), confirming the role of Panx1_EC_-mediated eATP in TRPV4_EC_ channel activation.

**Figure 2.**
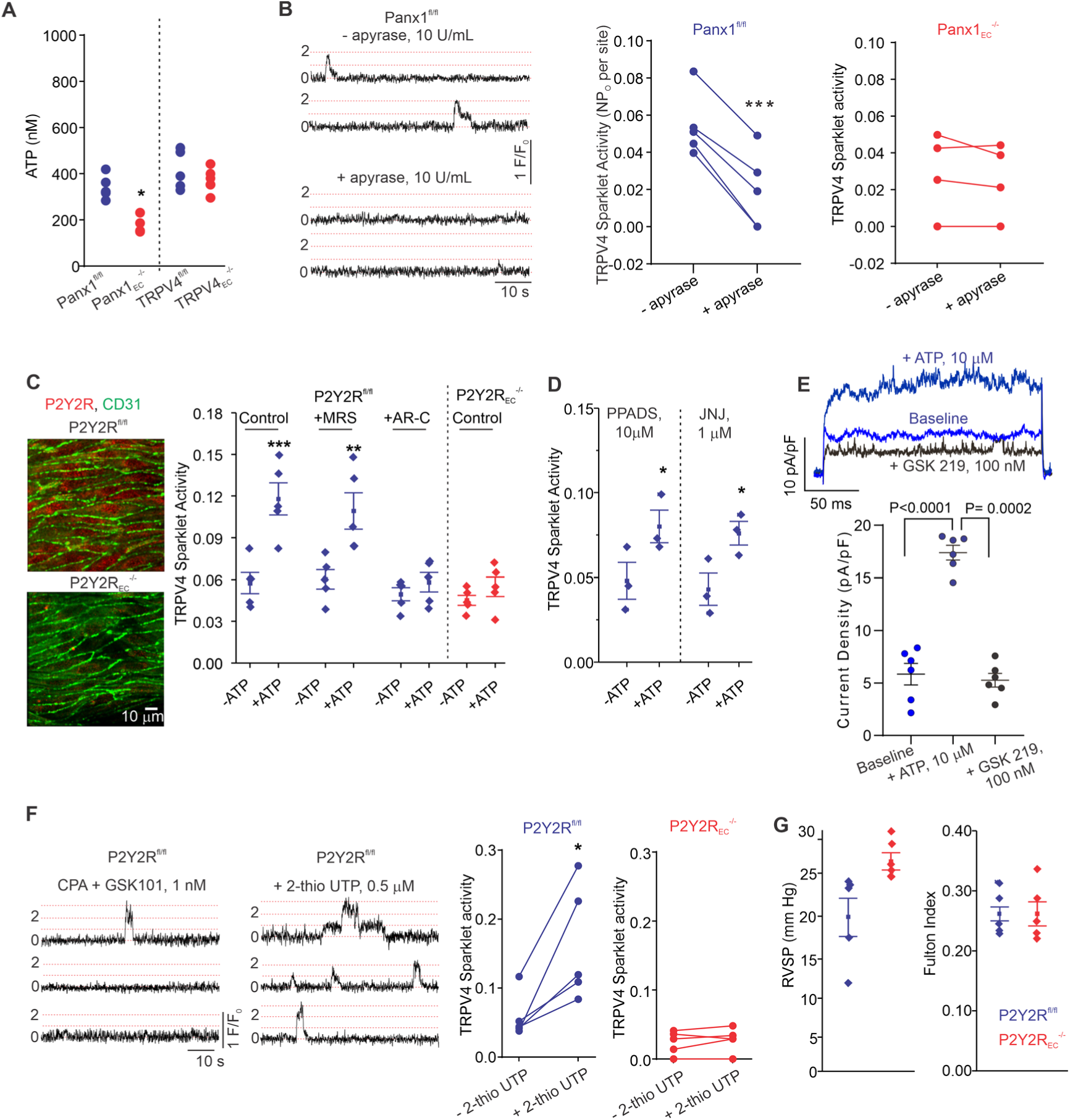
eATP activates TRPV4_EC_ channels via P2Y2R_EC_ stimulation. **A**, Release of ATP (nmol/L) from PAs of Panx1^fl/fl^, Panx1_EC_^-/-^, TRPV4^fl/fl^, and TRPV4_EC_^-/-^ mice (n = 5–6; \**P* < 0.05 vs. Panx1^fl/fl^; t-test]. **B,** *left,* Representative traces showing TRPV4_EC_ sparklet activity in *en face* preparations of PAs from Panx1^fl/fl^ mice in the absence or presence of apyrase (10 U/mL). Experiments were performed in Fluo-4–loaded PAs in the presence of CPA (20 μmol/L), included to eliminate Ca^2+^ release from intracellular stores. *Right,* TRPV4_EC_ sparklet activity (NP_O_) per site in *en face* preparations of PAs from Panx1^fl/fl^ and Panx1_EC_^-/-^ mice in the presence or absence of apyrase (10 U/mL; n = 5; \*\*\**P <* 0.001 vs. Panx1^fl/fl^ [-apyrase, 10 U/mL]; one-way ANOVA). **C,** *left,* Immunofluorescence images of *en face* 4^th^-order PAs from P2Y2R^fl/fl^ and P2Y2R_EC_^-/-^ mice*. Right,* Effects of ATP (10 μmol/L) on TRPV4_EC_ sparklet activity in the absence or presence of the P2Y1R inhibitor MRS2179 (MRS; 10 μmol/L) or P2Y2R inhibitor AR-C 118925XX (ARC; 10 μmol/L) in PAs from P2Y2R^fl/fl^ mice and P2Y2R_EC_^-/-^ mice, expressed as NP_O_ per site (n = 5; \*\*\**P <* 0.001, \*\**P <* 0.01 vs. -ATP; one-way ANOVA). **D,** Effects of ATP (10 μmol/L) on TRPV4_EC_ sparklet activity in the presence of the general P2X1-5/7R inhibitor PPADS (10 μmol/L) and P2X7R inhibitor JNJ-47965567 (JNJ; 1 μmol/L) in PAs of C57BL6/J mice (n = 5; \**P* < 0.05 vs. [-ATP, 10 μmol/L]; one-way ANOVA). **E,** *top,* representative ATP (10 μmol/L)–induced outward TRPV4 currents in freshly isolated ECs from C57BL6 mice and effect of GSK2193874 (GSK219, TRPV4 inhibitor, 100 nmol/L) in the presence of ATP, currents were elicited by a 200 ms voltage step from −50 mV to +100 mV; *bottom,* scatterplot showing outward currents at +100 mV under baseline conditions, after the addition of ATP, and after the addition of GSK219 (100 nM), n=6 cells, one-way ANOVA. **F,** *left,* Representative traces showing TRPV4_EC_ sparklet activity in *en face* preparations of PAs from P2Y2R^fl/fl^ mice in response to CPA (20 μmol/L) + GSK101 (1 nmol/L), CPA + 2-thio UTP (0.5 μmol/L), or CPA + GSK219 (100 nmol/L). *Right,* TRPV4_EC_ sparklet activity (NP_O_) per site in *en face* preparations of PAs from P2Y2R^fl/fl^ and P2Y2R_EC_^-/-^ mice under baseline conditions (i.e., 20 μmol/L CPA) and in response to 2-thio UTP (0.5 μmol/L; n = 5; \**P <* 0.05 vs. P2Y2R^fl/fl^ [-2-thio UTP]; two-way ANOVA). **G**, *left,* Average resting RVSP values in P2Y2R^fl/fl^ and P2Y2R_EC_^-/-^ mice (n = 6, \**P* < 0.05). *Right,* Average Fulton Index values in P2Y2R^fl/fl^ and P2Y2R_EC_^-/-^ mice (n = 5–6).

The pulmonary endothelium expresses both P2X and P2Y purinergic receptors^26–29^. The main P2Y receptor subtypes in the pulmonary endothelium are P2Y1R and P2Y2R^13, 26, 29^. The selective P2Y1R inhibitor MRS2179 (MRS, 10 μmol/L) did not alter eATP activation of TRPV4_EC_ sparklets (Fig. 2C). In contrast, the selective P2Y2R inhibitor AR-C 118925XX (AR-C; 10 μmol/L) completely abrogated the effect of eATP on TRPV4_EC_ sparklets (Fig. 2C). eATP was also unable to activate TRPV4_EC_ sparklets in inducible, endothelium-specific P2Y2R^-/-^ (P2Y2R_EC_^-/-^) mice (Fig. 2C), providing further evidence that eATP activates TRPV4_EC_ channels in PAs specifically via P2Y2R_EC_ signaling. The general P2X1-5 receptor inhibitor, PPADS (10 μmol/L), and P2X7 receptor inhibitor, JNJ-47965567 (JNJ, 1 μmol/L), did not alter the effect of eATP on TRPV4_EC_ sparklets, ruling out a role for P2X1-5/7 receptors in eATP activation of TRPV4_EC_ channels in PAs (Fig. 2D). In ECs freshly isolated from PAs of C57BL6 mice, ATP (10 μM) increased the outward currents through TRPV4 channels (Fig. 2E). Furthermore, the selective P2Y2R agonist, 2-thiouridine-5’-triphosphate (2-thio UTP; 0.5 μmol/L) activated TRPV4_EC_ sparklets in PAs from P2Y2R^fl/fl^ mice but not in PAs from P2Y2R_EC_^-/-^ mice (Fig. 2F).

Similar to TRPV4_EC_^-/-^ and Panx1_EC_^-/-^ mice, P2Y2R_EC_^-/-^ mice also showed elevated RVSP and an unaltered Fulton Index (Fig. 2G). Taken together, these findings demonstrate that P2Y2R_EC_ is the signaling intermediate for Panx1_EC_–TRPV4_EC_ interactions in PAs.

### Cav-1_EC_ provides a scaffold for Panx1_EC_**–**P2Y2R_EC_**–**TRPV4_EC_ signaling

Cav-1_EC_, an essential structural protein in the pulmonary circulation^21, 22, 30^, has been shown to co-localize with Panx1, P2Y2R, and TRPV4 channels in multiple cell types^17–19, 31^. Therefore, we hypothesized that Cav-1_EC_ provides a signaling scaffold that supports and maintains the spatial proximity among the individual elements in the Panx1_EC_–P2Y2R_EC_–TRPV4_EC_ pathway. To clearly delineate the role of Cav-1_EC_ in Panx1_EC_-dependent signaling, we utilized inducible, endothelium-specific Cav-1–knockout mice (Cav-1_EC_^-/-^; Fig. 3A and B). The loss of Cav-1_EC_ resulted in elevated RVSP in the absence of right ventricular hypertrophy (Fig. 3C), indicating a crucial role of Cav-1_EC_ in maintaining a low resting PAP. Baseline TRPV4_EC_ sparklet activity and activity induced by a low level of GSK101 (1 nmol/L) were reduced in PAs from Cav-1_EC_^-/-^ mice (Fig. 3D). However, higher-level activation of TRPV4_EC_ channels (30 nmol/L GSK101) resulted in similar TRPV4_EC_ sparklet activity between groups, suggesting that the number of functional TRPV4_EC_ channels is unaltered in Cav-1_EC_^-/-^ mice (Fig. 3D). Importantly, eATP-induced activation of TRPV4_EC_ sparklets was absent in PAs from Cav-1_EC_^-/-^ mice (Fig. 3E). These results provided the first functional evidence that Cav-1_EC_ is required for eATP–P2Y2R_EC_–TRPV4_EC_ signaling in PAs. To provide additional evidence to support Cav-1_EC_–dependent co-localization of Panx1_EC_–P2Y2R_EC_– TRPV4_EC_ signaling elements in PAs, we performed *in situ* proximity ligation assays (PLA), which confirmed that Cav-1_EC_ exists within nanometer proximity of Panx1_EC_, P2Y2R_EC_, and TRPV4_EC_ channels in PAs (Fig. 3F).

**Figure 3.**
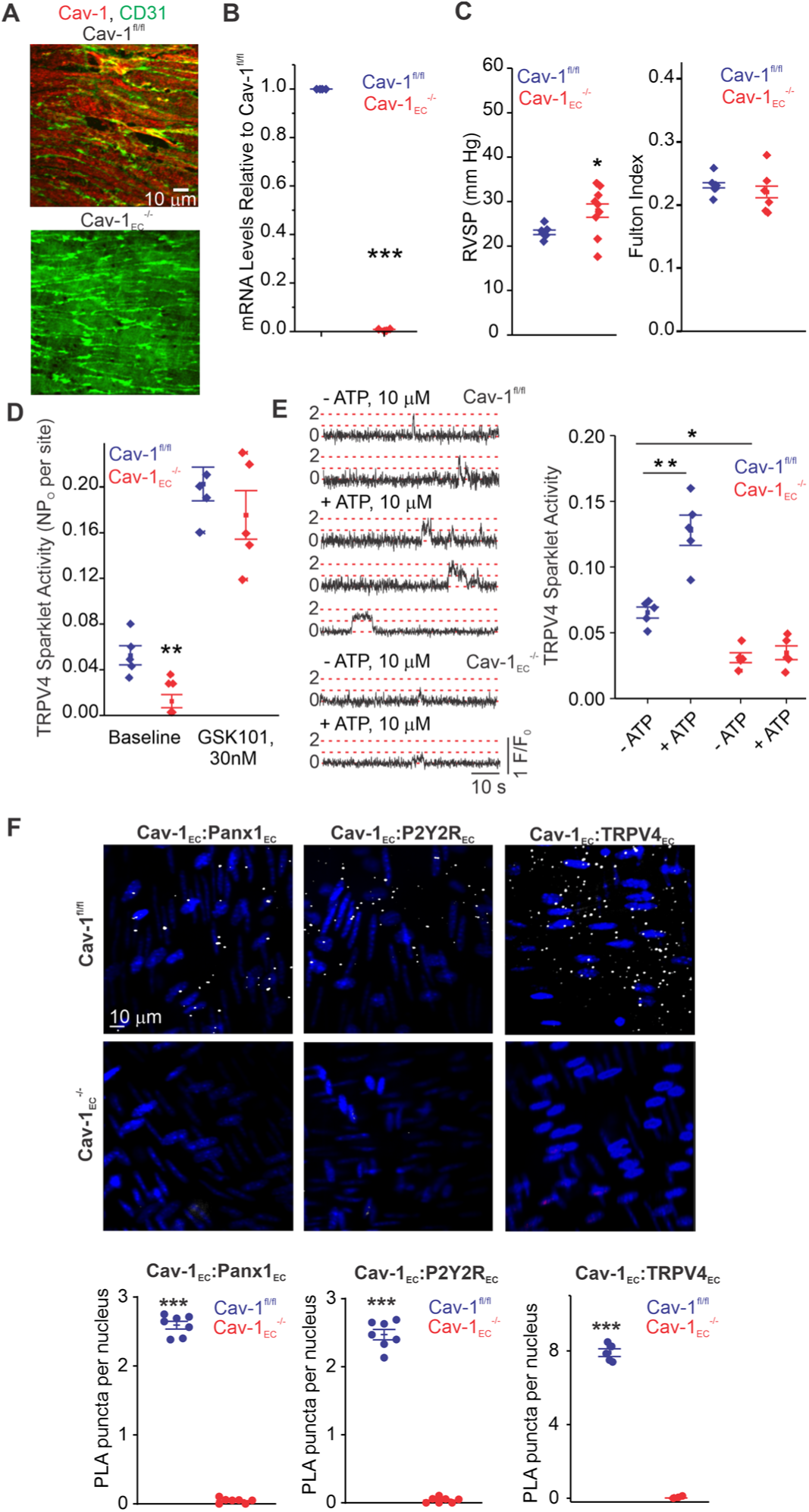
Cav-1_EC_ provides a signaling scaffold for Panx1_EC_–P2Y2R_EC_–TRPV4_EC_ signaling in PAs. **A**, Immunofluorescence images of *en face* 4^th^-order PAs from Cav-1^fl/fl^ (*top*) and Cav-1_EC_^-/-^ (*bottom*) mice. **B,** Endothelial Cav-1 mRNA levels in PAs relative to those in Cav-1^fl/fl^ mice (n = 4; \*\*\**P <* 0.001; t-test). **C,** *left,* Average resting RVSP values in Cav-1^fl/fl^ and Cav-1_EC_^-/-^ mice (n = 6; \**P <* 0.05 vs. Cav-1^fl/fl^; one-way ANOVA). *Right,* Average Fulton Index values in Cav-1^fl/fl^ and Cav-1_EC_^-/-^ mice (n = 5–6). **D**, TRPV4_EC_ sparklet activity (NP_O_) per site in *en face* preparations of PAs from Cav-1^fl/fl^ and Cav-1_EC_^-/-^ mice at baseline and in response to 30 nmol/L GGSK101 (n = 6; \*\**P* < 0.01 vs. baseline Cav-1^fl/fl^; one-way ANOVA). Experiments were performed in Fluo-4–loaded 4^th^-order PAs in the presence of CPA (20 μmol/L), included to eliminate Ca^2+^ release from intracellular stores. **E***, left,* Representative traces showing TRPV4_EC_ sparklets in *en face* preparations of PAs from Cav-1^fl/fl^ and Cav-1_EC_^-/-^ mice in the presence and absence of ATP (10 μmol/L). *Right,* TRPV4_EC_ sparklet activity (NP_O_) per site in *en face* preparations of PAs from Cav-1^fl/fl^ and Cav-1_EC_^-/-^ mice in the presence or absence of 10 μmol/L ATP (n = 5; \**P <* 0.05; \*\**P* < 0.01 vs. [-ATP] Cav-1^fl/fl^; two-way ANOVA). **F,** *top,* Representative merged images of proximity ligation assays (PLA) showing EC nuclei and Cav-1_EC_:Panx1_EC_, Cav-1_EC_:P2Y2R_EC_, and Cav-1_EC_:TRPV4_EC_ co-localization (white puncta) in 4^th^-order PAs from Cav-1^fl/fl^ and Cav-1_EC_^-/-^ mice. *Bottom,* Quantification of Cav-1_EC_:Panx1_EC_, Cav-1_EC_:P2Y2R_EC_, and Cav-1_EC_:TRPV4_EC_ co-localization in PAs from Cav-1^fl/fl^ and Cav-1_EC_^-/-^ mice (n = 5; *** *P* < 0.001 vs. Cav-1^fl/fl^; t-test).

### Cav-1_EC_ anchoring of PKCα mediates P2Y2R_EC_-dependent activation of TRPV4_EC_ channels in PAs

P2Y2R is a Gq protein-coupled receptor that activates the phospholipase C (PLC)– diacylglycerol (DAG)–PKC signaling pathway. Notably, PKC is known to phosphorylate TRPV4 channels and potentiate its activity^32^. eATP, the DAG analog OAG (1 μmol/L), and the PKC activator phorbol myristate acetate (PMA; 10 nmol/L) stimulated TRPV4_EC_ sparklet activity in small PAs (Fig. 4A–C). Inhibition of PLC with U73122 (3 μmol/L) abolished eATP activation of TRPV4_EC_ sparklets, but not OAG- or PMA-induced activation of TRPV4_EC_ sparklets. Moreover, the PKC*α/β* inhibitor Gö-6976 (1 μmol/L) prevented activation of TRPV4_EC_ sparklets by ATP, OAG and PMA (Fig. 4A–C), supporting the concept that eATP activation of P2Y2R_EC_ stimulates TRPV4_EC_ channel activity via PLC–DAG–PKC signaling in PAs. TRPV4_EC_ channel activation by PLC–DAG–PKC signaling was further supported by increased activity of TRPV4_EC_ sparklets in PAs from Cdh5-optoα1 adrenergic receptor (Cdh5-optoα1AR) mouse, which expresses light-sensitive α1AR in endothelial cells. When activated with light (∼473 nm), Optoα1AR generates the secondary messengers IP3 and diacylglycerol (DAG)^33^. Light activation resulted in increased activity of TRPV4_EC_ sparklets (Fig. 4D), an effect that was abolished by the PKCα/β inhibitor Gö-6976 (1 μM).

**Figure 4.**
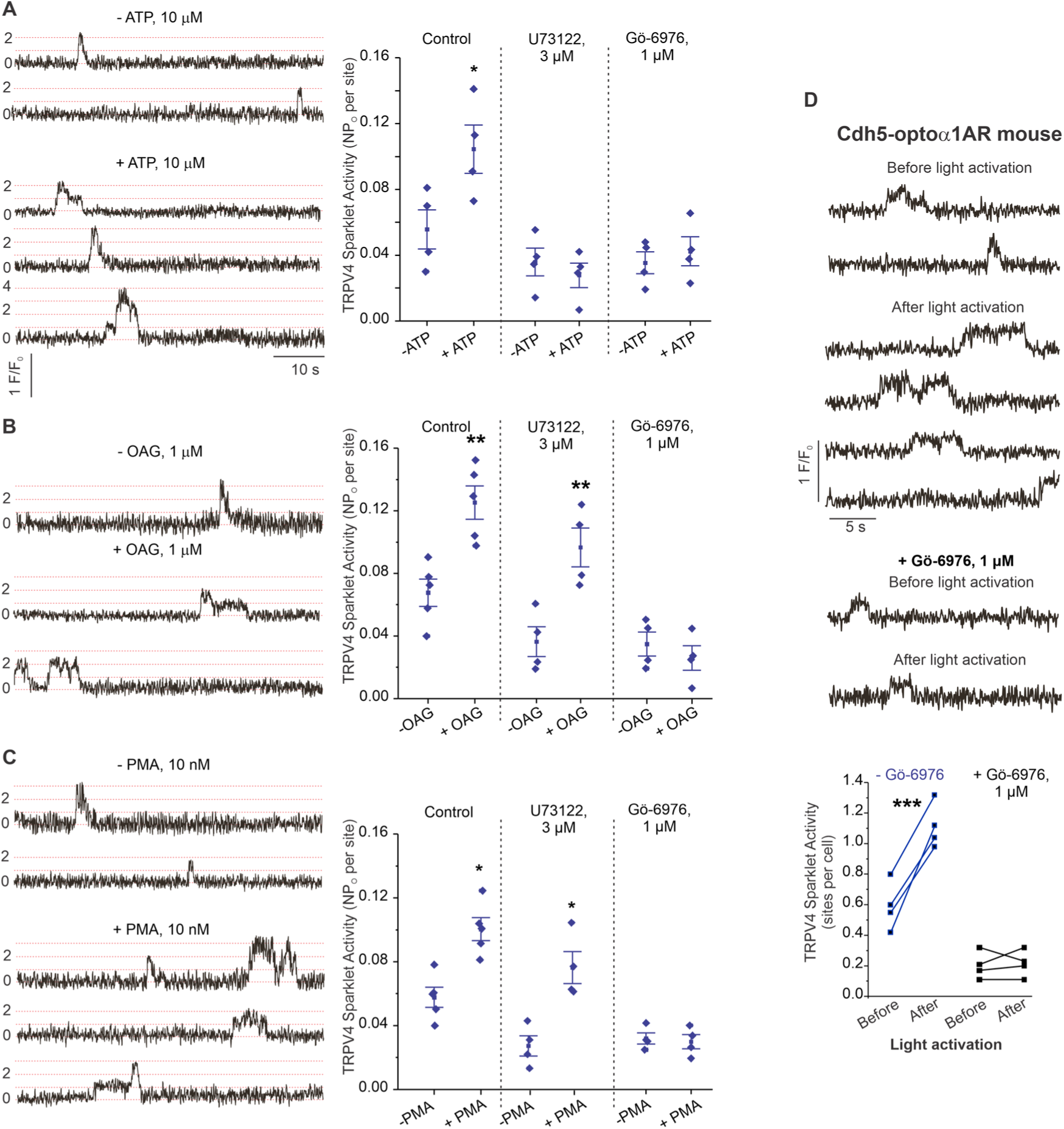
eATP activates TRPV4_EC_ channels via P2Y2R_EC_–PLC–PKC signaling in PAs. **A**, *left,* Representative traces showing TRPV4_EC_ sparklet activity in *en face* preparations of PAs from C57BL6/J mice before and after treatment with ATP (10 μmol/L). *Right,* Effects of U73122 (PLC inhibitor; 3 μmol/L) or Gö-6976 (PKCα/β inhibitor; 1 μmol/L) on TRPV4_EC_ sparklet activity in *en face* preparations of PAs from C57BL6/J mice before and after treatment with ATP (10 μmol/L), expressed as NP_O_ per site. Experiments were performed in Fluo-4–loaded 4^th^-order Pas in the presence of CPA (20 μmol/L), included to eliminate Ca^2+^ release from intracellular stores (n = 5; \**P* < 0.05 vs. [-ATP]; one-way ANOVA). **B,** *left,* Representative traces showing TRPV4_EC_ sparklet activity in *en face* preparations of PAs from C57BL6/J mice in the absence or presence of OAG (DAG analogue; 1 μmol/L). *Right,* Effects of U73122 (3 μmol/L) or Gö-6976 (1 μmol/L) on TRPV4_EC_ sparklet activity in *en face* preparations of PAs from C57BL6/J mice before and after treatment with OAG (1 μmol/L), expressed as NP_O_ per site (n = 6; \*\**P* < 0.01 vs. [-OAG]; one-way ANOVA). **C,** *left,* Representative traces showing TRPV4_EC_ sparklets in *en face* preparations of PAs from C57BL6/J mice in the absence or presence of PMA (PKC activator; 10 nmol/L). *Right,* Effects of U73122 (3 μmol/L) or Gö-6976 (1 μmol/L) on TRPV4_EC_ sparklet activity in *en face* preparations of PAs from C57BL6/J mice before and after treatment with PMA (10 nmol/L), expressed as NP_O_ per site (n = 6; \**P* < 0.05 vs. [-PMA]; one-way ANOVA). **D.** *top,* Representative traces showing TRPV4_EC_ sparklet activity in *en face* preparations of PAs from Cdh5-optoα1AR (adrenergic receptor) mouse before and after the light activation (470 nm); *bottom*, scatter plot showing the sparklet activity, expressed as sparklet sites per cell, before and after the light activation, in the absence or presence of PKCα/β inhibitor Gö-6976 (1 μM, n=4, \*\*\**P* < 0.001).

Since Cav-1 possesses a PKC-binding domain^34^ and exists in nanometer proximity with TRPV4_EC_ channels and P2Y2R_EC_, we tested the hypothesis that Cav-1_EC_ anchoring of PKC mediates P2Y2R_EC_–TRPV4_EC_ channel interaction in PAs. PLA experiments confirmed that PKC also exists in nanometer proximity with Cav-1_EC_ in PAs (Fig. 5A). The PKC**-**dependence of Cav-1_EC_ activation of TRPV4_EC_ channels was confirmed by studies in HEK293 cells transfected with TRPV4 alone or TRPV4 channels plus Cav-1 (Fig. 5B), which showed that TRPV4 currents were increased in the presence of Cav-1. Further, the PKC*α/β* inhibitor Gö-6916 (1 μmol/L) reduced TRPV4 channel currents in Cav-1/TRPV4–co-transfected cells to the level of that in cells transfected with TRPV4 alone (Fig. 5B and C). These results imply that Cav-1 enhances TRPV4 channel activity via PKC*α/β* anchoring. Experiments in which TRPV4 channels were co-expressed with PKCα or PKCβ showed that only PKCα increased currents through TRPV4 channels (Fig. 5D). Collectively, these results support the conclusion that Panx1_EC_–P2Y2R_EC_– PKCα–TRPV4_EC_ signaling on a Cav-1_EC_ scaffold reduces PA contractility and lowers resting PAP (Fig. 5E).

**Figure 5.**
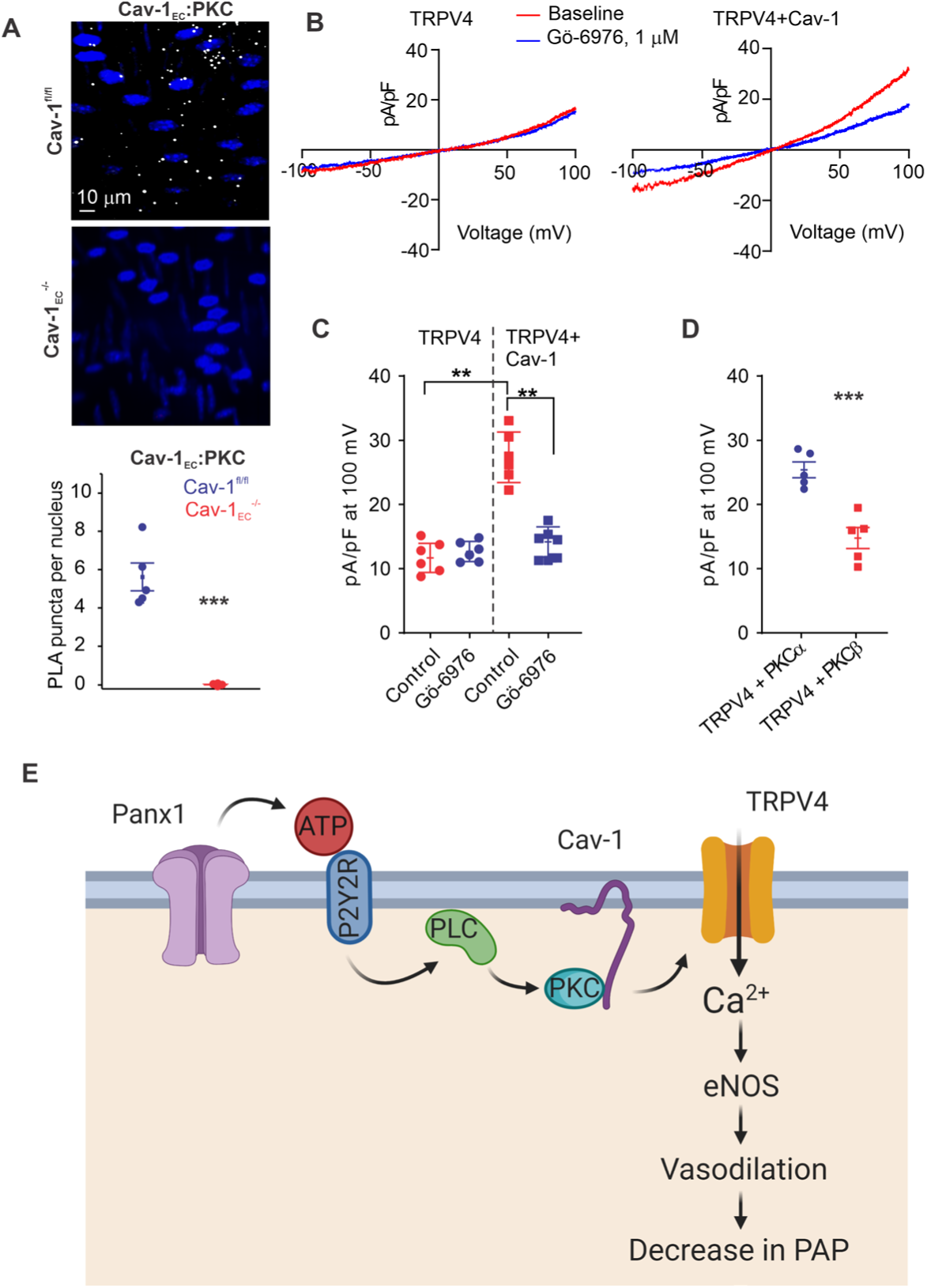
Localization of PKCα with Cav-1_EC_ increases the activity of TRPV4_EC_ channels. **A**, *top,* Representative merged images of proximity ligation assays (PLA) showing EC nuclei and Cav-1:PKC co-localization (white puncta) in 4^th^-order PAs from Cav-1^fl/fl^ and Cav-1_EC_^-/-^ mice. *Bottom,* Quantification of Cav-1:PKC co-localization in PAs from Cav-1^fl/fl^ and Cav-1_EC_^-/-^ mice (n = 5; \*\*\**P* < 0.001 vs. Cav-1^fl/fl^; t-test). **B,** Representative traces showing TRPV4 currents in the absence or presence of Gö-6976 (PKC inhibitor; 1 μmol/L) in HEK293 cells transfected with TRPV4 only or co-transfected with TRPV4 plus wild-type Cav-1, recorded in the whole-cell patch-clamp configuration. **C,** Current density plot of TRPV4 currents at +100 mV in the absence or presence of Gö6976 (1 μmol/L) in HEK293 cells transfected with TRPV4 or TRPV4 + Cav-1 (n = 5; \*\**P <* 0.01 vs. Control; one-way ANOVA). **D,** Current density plot of TRPV4 currents at +100 mV in HEK293 cells transfected with TRPV4 + PKCα or TRPV4 + PKCβ (n = 5; \*\*\**P* < 0.001 vs. TRPV4 + PKCα; t-test). **E,** Schematic depiction of the Panx1_EC_–P2Y2R_EC_–TRPV4_EC_ signaling pathway that promotes vasodilation and lowers PAP in PAs. ATP released from Panx1_EC_ channels activates P2Y2R_EC_ purinergic receptors on the EC membrane. Stimulation of P2Y2R_EC_ recruits PKCα, which anchors to the scaffolding protein Cav-1_EC_ in close proximity to TRPV4_EC_ channels. TRPV4_EC_ channel-dependent vasodilation lowers pulmonary arterial pressure (PAP).

## Discussion

Regulation of PA contractility and PAP is a complex process involving multiple cell types and signaling elements. In particular, the endothelial signaling mechanisms that control resting PAP remain poorly understood. Our studies identify a Panx1_EC_ and TRPV4_EC_ channel-containing signaling nanodomain that reduces PA contractility and lowers PAP. Although both Panx1_EC_ and TRPV4_EC_ channels have been implicated in dilation of systemic arteries, their impact on PAP remains unknown. We demonstrate critical roles for several key, linked mechanistic, pathways showing that 1) Panx1_EC_ increases eATP levels in small PAs; 2) Panx1_EC_-generated eATP, in turn, enhances Ca^2+^ influx through TRPV4_EC_ channels, thereby dilating PAs and lowering PAP; 3) eATP acts through purinergic P2Y2R_EC_–PKCα signaling to activate TRPV4_EC_ channels; and 4) Cav-1_EC_ provides a signaling scaffold that ensures spatial proximity among the elements of the Panx1_EC_–P2Y2R_EC_–PKCα–TRPV4_EC_ pathway. Our findings reveal a novel signaling axis that can be engaged by physiological stimuli to lower PAP and could also be therapeutically targeted in pulmonary vascular disorders. Moreover, the conclusions in this study may assist in future investigations of the mechanisms underlying pulmonary endothelial dysfunction.

Both ECs and SMCs control vascular contractility and arterial pressure. The expression of Panx1 and TRPV4 channels in both ECs and SMCs^8, 18, 35–37^ makes it challenging to decipher the cell type-specific roles of Panx1 and TRPV4 channels using global knockouts or pharmacological strategies. Therefore, studies utilizing EC- or SMC-specific knockout mice are necessary for a definitive assessment of the control of PAP by EC and SMC Panx1 and TRPV4 channels. Although SMC TRPV4 channels have been shown to contribute to hypoxia-induced pulmonary vasoconstriction, resting PAP is not altered in global TRPV4^-/-^ mice^38, 39^. Further, our studies indicate that SMC Panx1 and TRPV4 channels do not influence resting PAP. Taken together with findings from EC-knockout mice, these results provide strong evidence that endothelial, but not SMC, Panx1 and TRPV4 channels maintain low PA contractility and PAP under basal conditions.

Recent studies in pulmonary fibroblasts and other cell types suggest that TRPV4 channel-mediated increases in cytosolic Ca^2+^ can induce eATP release through Panx1^40, 41^. However, the reverse interaction, in which Panx1-mediated eATP release activates TRPV4 channels, has not been explored in any cell type. Since Panx1 is activated by cytosolic Ca^2+ 42^ and eATP has been previously shown to activate TRPV4_EC_ channels^9^, bidirectional signaling between Panx1 and TRPV4 channels is conceivable. However, our demonstration that baseline eATP levels are unchanged in PAs from TRPV4_EC_^-/-^ mice rules out a role for TRPV4_EC_ channels in controlling eATP release under baseline conditions. Nevertheless, these data from pulmonary ECs do not rule out potential TRPV4–Ca^2+^–Panx1 signaling in other cell types.

Elevated capillary TRPV4_EC_ channel activity has been linked to increased endothelial permeability^43, 44^, lung injury^45^, and pulmonary edema^43, 44^. Moreover, Panx1_EC_-mediated eATP release is associated with vascular inflammation at the level of capillaries^8^. The physiological roles of Panx1_EC_ and TRPV4_EC_ channels in PAs, however, remain unknown. ECs from pulmonary capillaries and arteries are structurally and functionally different. Whereas PAs control pulmonary vascular resistance and PAP, capillaries control vascular permeability. TRPV4_EC_ channels couple with distinct targets in arterial and capillary ECs ^25, 46^. Our data identify physiological roles of Panx1_EC_–TRPV4_EC_ channel signaling in PAs, but whether such signaling operates in the capillary endothelium and is essential for its physiological function is unclear.

Purinergic signaling and the endogenous purinergic receptor agonist eATP are essential controllers of pulmonary vascular function^13, 26, 28, 47^. Our discovery of the Panx1_EC_–P2Y2R_EC_– TRPV4_EC_ pathway establishes a signaling axis in ECs that regulates pulmonary vascular function. The pulmonary vasculature is a high-flow circulation, and pulmonary ECs have been shown to release eATP in response to flow/shear stress^12^. Therefore, flow/shear stress could be a potential physiological activator of Panx1_EC_–P2Y2R_EC_–TRPV4_EC_ signaling in PAs. Further studies are needed to verify this possibility. Several purinergic receptor subtypes are expressed in the pulmonary vasculature, including P2YRs and P2XRs^26–28^. Although only P2Y2R_EC_ appears to mediate eATP activation of TRPV4_EC_ channels, our studies do not rule out potentially important roles for other P2Y or P2X receptors in the pulmonary endothelium.

Activation of TRPV4_EC_ channels by eATP released through Panx1_EC_ in PAs would be facilitated by spatial localization of TRPV4_EC_ channels with Panx1_EC_. In keeping with this, several scaffolding proteins are known to promote localization of TRPV4 channels with their regulatory proteins, including A-kinase anchoring protein 150 (AKAP150) and Cav-1^23, 48^. Although AKAP150 is not found in the pulmonary endothelium^9^, Cav-1 is a key structural protein in the pulmonary vasculature and has a well-established role in controlling pulmonary vascular function, as demonstrated by increased RVSP in global Cav-1^-/-^ mice^49, 50^. Moreover, Cav-1–dependent signaling is impaired in pulmonary hypertension^20–22^. Studies in other cell types have shown that Cav-1 can co-localize with Panx1 and P2Y2Rs^18, 19^. Additionally, Cav-1 can interact with PKC at the Cav-1 scaffolding domain^34^. Our results demonstrate that Cav-1_EC_ exists in nanometer proximity with Panx1_EC_, P2Y2R_EC_, PKC, and TRPV4_EC_ channels in PAs. Furthermore, the activation of TRPV4_EC_ channels by Panx1_EC_, eATP, P2Y2R_EC_ or PKCα requires Cav-1_EC_. Based on these findings, we conclude that Cav-1_EC_ enables Panx1_EC_–P2Y2R_EC_–TRPV4_EC_ signaling at EC membranes in PAs. Cav-1 is also a well-known anchor protein for eNOS^16^, acting by stabilizing eNOS expression and negatively regulating its activity^16^. We previously showed that TRPV4_EC_ Ca^2+^ sparklets activate eNOS in PAs^9, 36^. Thus, Cav-1_EC_ enhancement of Ca^2+^ influx through TRPV4_EC_ channels may represent novel mechanisms for regulating eNOS activity.

Cav-1_EC_/PKCα-dependent signaling is a novel endogenous mechanism for activating arterial TRPV4_EC_ channels and lowering PAP. Proximity to PKCα appears to be crucial for the normal function of TRPV4 channels. Evidence from the systemic circulation suggests that co-localization of TRPV4 channels with scaffolding proteins enhances their activity^51, 52^, and we specifically demonstrated that PKC anchoring by AKAP150 enhances the activity of TRPV4_EC_ channels in mesenteric arteries^23^. Here, we show that PKC anchoring by Cav-1_EC_ enables PKC activation of TRPV4_EC_ channels in PAs. This discovery raises the possibility that disruption of PKC anchoring by Cav-1_EC_ could impair the Panx1_EC_–P2Y2R_EC_–TRPV4_EC_ signaling axis under disease conditions. A lack of PKC anchoring by scaffolding proteins in systemic arteries has been demonstrated in obesity and hypertension^23, 52^. Further studies of pulmonary vascular disorders are required to establish whether the Panx1_EC_–P2Y2R_EC_–PKCα–TRPV4_EC_ signaling axis is impaired in pulmonary vascular disorders.

In conclusion, Panx1_EC_–TRPV4_EC_ signaling reduces PA contractility and maintains a low resting PAP. This mechanism is facilitated by eATP released through Panx1_EC_ and subsequent activation of P2Y2R_EC_–PKCα signaling. Cav-1_EC_ ensures the spatial proximity among Panx1_EC_, P2Y2R_EC_, and TRPV4_EC_ channels and also anchors PKCα close to TRPV4_EC_ channels. These findings identify a novel endothelial Ca^2+^ signaling mechanism that reduces PA contractility. Further investigations are needed to determine whether impairment of this pathway contributes to elevated PAP in pulmonary vascular disorders and whether this pathway can be targeted for therapeutic benefit.

## Materials and Methods

### Drugs and chemical compounds

Cyclopiazonic acid (CPA), GSK2193874, GSK1016790A, Phorbol 12-myristate 13-acetate (PMA), AR-C 118925XX, 2-Thio UTP tetrasodium salt, MRS2179, U-73122 and NS309 were purchased from Tocris Bioscience (Minneapolis, MN, USA). Fluo-4-AM (Ca^2+^ indicator) were purchased from Invitrogen (Carlsbad, CA, USA). 1-O-9Z-octadecenoyl-2-O-acetyl-*sn*-glycerol (OAG), PPADS (sodium salt), Gö-6976, JNJ-47965567 and U46619 were purchased from Cayman Chemicals (Ann Arbor, MI, USA). Tamoxifen and apyrase were obtained from Sigma-Aldrich (St. Louis, MO, USA).

### Animal protocols and models

All animal protocols were approved by the University of Virginia Animal Care and Use Committee (protocols 4100 and 4120). This study was performed in strict accordance with the recommendations in the Guide for the Care and Use of Laboratory Animals of the National Institutes of Health. For surgical procedures, every effort was made to minimize suffering. Both male and female mice were used in this study and age- and sex-matched controls were used. C57BL6/J were obtained from the Jackson Laboratory (Bar Harbor, ME). Inducible endothelial cell (EC)-specific TRPV4 channel knockout (TRPV4_EC_^-/-^)^53, 54^, smooth muscle cell (SMC)-specific TRPV4 channel knockout (TRPV4_SMC_^-/-^)^55^, EC caveolin-1 knockout (Cav-1_EC_^-/-^)^56^, EC-specific P2Y2R receptor knockout (P2Y2R_EC_^-/-^)^57^, EC-specific Panx1 channel knockout (Panx1_EC_^-/-^)^24, 54^ and SMC-specific Panx1 channel knockout (Panx1_SMC_^-/-^)^55^ mice (10-14 weeks old) were used. The mouse strain Cdh5-optoα1AR was developed by CHROMus^TM^ which is supported by the National Heart Lung Blood Institute of the National Institute of Health under award number R24HL120847. Mice were housed in an enriched environment and maintained under a 12:12 h light/dark photocycle at ∼23°C with fresh tap water and standard chow diet available *ad libitum.* Mice were euthanized with pentobarbital (90 mg/kg^-1^; intraperitoneally; Diamondback Drugs, Scottsdale, AZ) followed by cervical dislocation for harvesting lung tissue. Fourth-order pulmonary arteries (PAs, ∼50 μm diameter) were isolated in cold HEPES-buffered physiological salt solution (HEPES-PSS, in mmol/L, 10 HEPES, 134 NaCl, 6 KCl, 1 MgCl_2_ hexahydrate, 2 CaCl_2_ dihydrate, and 7 dextrose, pH adjusted to 7.4 using 1 mol/L NaOH).

TRPV4^fl/fl 53^, Cav-1^fl/fl 56^, Panx1^fl/fl 24, 54^ and P2Y2R^fl/fl 57^mice were crossed with VE-Cadherin (Cdh5, endothelial) Cre mice^53^ or SMMHC (smooth muscle) Cre mice ^58^. EC- or SMC-specific knockout of TRPV4, Cav-1, Panx1, or P2Y2R was induced by injecting 6 week-old TRPV4^fl/fl^ Cre^+^, Cav-1^fl/fl^ Cre^+^, Panx1^fl/fl^ Cre^+^ and P2Y2R^fl/fl^ Cre^+^ mice with tamoxifen (40 mg/kg intraperitoneally per day for 10 days). Tamoxifen-injected TRPV4^fl/fl^ Cre^-^, Cav-1^fl/fl^ Cre^-^, Panx1^fl/fl^ Cre^-^ and P2Y2R^fl/fl^ Cre^-^ mice were used as controls. Mice were used for experiments after a two-week washout period. Genotypes for Cdh5 Cre and SMMHC Cre were confirmed following previously published protocols^53, 58^. TRPV4^fl/fl53^, Cav-1^fl/fl56^, Panx1^fl/fl24, 54^, P2Y2R^fl/fl57^ genotyping was performed as described previously. Cdh5-Optoα1AR mice were developed by CHROMus (Cornell University, USA).

### Right ventricular systolic pressure (RVSP) and Fulton Index measurement

Mice were anesthetized with pentobarbital (50 mg/kg bodyweight; intraperitoneally) and bupivacaine HCl (100 μL of 0.25% solution; subcutaneously) was used to numb the dissection site on the mouse. RVSP was measured as an indirect indicator of pulmonary arterial pressure (PAP). A Mikro-Tip pressure catheter (SPR-671; Millar Instruments, Huston, TX), connected to a bridge amp (FE221), and a PowerLab 4/35 4-channel recorder (Instruments, Colorado Springs, CO), was cannulated through the external jugular vein into the right ventricle. Right ventricular pressure and heart rate were acquired and analyzed using LabChart8 software (ADInstruments, Colorado Springs, CO). A stable 3-minute recording was acquired for all the animals, and 1-minute continuous segment was used for data analysis. When necessary, traces were digitally filtered using a low-pass filter at a cut-off frequency of 50 Hz. At the end of the experiments, mice were euthanized, and the hearts were isolated for right ventricular hypertrophy analysis. Right ventricular hypertrophy was determined by calculating the Fulton Index, a ratio of the right ventricular (RV) heart weight over the left ventricular (LV) plus septum (S) weight (RV/ LV+S).

### Luciferase assay for total ATP release

ATP assay protocol was adapted from Yang et al.^59^. Fourth-order pulmonary arteries (PAs, ∼ 50 μm diameter) were isolated in cold HEPES-buffered physiological salt solution (HEPES-PSS, in mmol/L, 10 HEPES, 134 NaCl, 6 KCl, 1 M gCl _2_ hexahydrate, 2 CaCl_2_ dihydrate, and 7 dextrose, pH adjusted to 7.4 using 1 mol/L NaOH). Isolated PAs were pinned down *en face* on a Sylgard block and cut open. PAs were placed in black, opaque 96-well plates and incubated in HEPES-PSS for 10 minutes at 37 °C, followed by incubation with the ectonucleotidase inhibitor ARL 67156 (300 μmol/L, Tocris Bioscience, Minneapolis, MN) for 30 minutes at 37 °C. 50 μL volume of each sample was transferred to another black, opaque 96-well plate. ATP was measured using ATP bioluminescence assay reagent ATP Bioluminescence HSII kit (Roche Applied Science, Penzberg, Germany). Using a luminometer (FluoStar Omega), 50 μL of luciferin:luciferase reagent (ATP bioluminescence assay kit HSII; Roche Applied Science, Penzberg, Germany) was injected into each well and luminescence was recorded following a 5 second orbital mix and sample measurement at 7 seconds. ATP concentration in each sample was calculated from an ATP standard curve.

### Cardiac Magnetic Resonance Imaging (MRI)

MRI studies were conducted under protocols that comply with the Guide for the Care and Use of Laboratory Animals (NIH publication no. 85-23, Revised 1996). Mice were positioned in the scanner under 1.25% isoflurane anesthesia and body temperature was maintained at 37°C using thermostatic circulating water. A cylindrical birdcage RF coil (30 mm-diameter, Bruker, Ettlingen, Germany) with an active length of 70 mm was used, and heart rate, respiration, and temperature were monitored during imaging using a fiber optic, MR-compatible system (Small Animal Imaging Inc., Stony Brook, NY). MRI was performed on a 7 Tesla (T) Clinscan system (Bruker, Ettlingen, Germany) equipped with actively shielded gradients with a full strength of 650 mT/m and a slew rate of 6666 mT/m/ms^60^. Six short-axis slices were acquired from base to apex, with slice thickness of 1 mm, in-plane spatial resolution of 0.2 × 0.2 mm^2^, and temporal resolution of 8–12 ms. Baseline ejection fraction (EF), end-diastolic volume (EDV), end-systolic volume (ESV), myocardial mass, wall thickness, stroke volume (SV), and cardiac output (CO) were assessed from the cine images using the freely available software Segment version 2.0 R5292 (http://segment.heiberg.se).

### Pressure myography

Isolated mouse PAs (∼ 50 μm) were cannulated on glass micropipettes in a pressure myography chamber (The Instrumentation and Model Facility, University of Vermont, Burlington, VT) at areas lacking branching points, and were pressurized at a physiological pressure of 15 mm Hg^36^. Arteries were superfused with PSS (in mmol/L, 119 NaCl, 4.7 KCl, 1.2 KH_2_PO_4_, 1.2 MgCl_2_ hexahydrate, 2.5 CaCl_2_ dihydrate, 7 dextrose, and 24 NaHCO_3_) at 37°C and bubbled with 20% O_2_/5% CO_2_ to maintain the pH at 7.4. All drug treatments were added to the superfusing PSS. PAs were pre-constricted with 50 nmol/L U46619 (a thromboxane A2 receptor agonist). All other pharmacological treatments were performed in the presence of U46619. Before measurement of vascular reactivity, arteries were treated with NS309 (1 μmol/L), a direct opener of endothelial IK/SK channels, to assess endothelial health. Arteries that failed to fully dilate to NS309 were discarded. Changes in arterial diameter were recorded at a 60-ms frame rate using a charge-coupled device camera and edge-detection software (IonOptix LLC, W estwood, M A) ^25, 52^. All drug treatments were incubated for 10 minutes. At the end of each experiment, Ca^2+^-free PSS (in mmol/L, 119 NaCl, 4.7 KCl, 1.2 KH_2_PO_4_, 1.2 MgCl_2_ hexahydrate, 7 dextrose, 24 NaHCO_3_, and 5 EGTA) was applied to assess the maximum passive diameter. Percent constriction was calculated by:

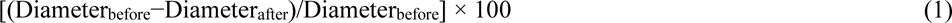

where Diameter_before_ is the diameter of the artery before a treatment and Diameter_after_ is the diameter after the treatment. Percent dilation was calculated by:

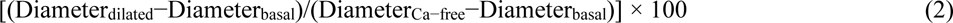

where Diameter_basal_ is the stable diameter before drug treatment, Diameter_dilated_ is the diameter after drug treatment, and Diameter_Ca-free_ is the maximum passive diameter.

### Ca^2+^ imaging

Measurements of TRPV4_EC_ Ca^2+^ sparklets in the native endothelium of mouse PAs were performed as previously described^25^. Briefly, 4^th^-order (∼ 50 μm) PAs were pinned down *en face* on a Sylgard block and loaded with fluo-4-AM (10 μmol/L) in the presence of pluronic acid (0.04%) at 30°C for 30 minutes. TRPV4_EC_ Ca^2+^ sparklets were recorded at 30 frames per second with Andor Revolution WD (with Borealis) spinning-disk confocal imaging system (Oxford Instruments, Abingdon, UK) comprised of an upright Nikon microscope with a 60X water dipping objective (numerical aperture 1.0) and an electron multiplying charge coupled device camera (iXon 888, Oxford Instruments, Abingdon, UK). All experiments were carried out in the presence of cyclopiazonic acid (20 μmol/L, a sarco-endoplasmic reticulum (ER) Ca ^2+^-ATPase inhibitor) in order to eliminate the interference from Ca^2+^ release from intracellular stores. Fluo-4 was excited at 488 nm with a solid-state laser and emitted fluorescence was captured using a 525/36-nm band-pass filter. TRPV4_EC_ Ca^2+^ sparklets were recorded before and 5 minutes after the addition of specific compounds. To generate fractional fluorescence (F/F_0_) traces, a region of interest defined by a 1.7-μm ^2^ (5×5 pixels) box was placed at a point corresponding to peak sparklet amplitude. Each field of view was ∼ 110×110 μm and covered ∼ 15 ECs. Representative F/F_0_ traces were filtered using a Gaussian filter and a cutoff corner frequency of 4 Hz. Sparklet activity was assessed as described previously using the custom-designed SparkAn software ^25, 52^.

For the experiments in Cdh5-optoα1AR mice, PAs were loaded with X-Rhod-1 AM (5 μM, Thermo Fisher Scientific Inc., Waltham, MA, USA) for 30 minutes at 30°C. X-Rhod-1 was excited at 561 nm and the emitted light was captured with a 607/36-nm band-pass filter. Optoα1AR was activated at 470 nm for 5 seconds using pE-4000 (CoolLED Ltd, Andover, UK).

### Calculation of TRPV4 sparklet activity per site

Activity of TRPV4 Ca^2+^ sparklets was evaluated as described previously^25, 52^. Area under the curve for all the events at a site was determined using trapezoidal numerical integration ([F−F_0_]/F_0_ over time, in seconds). The average number of active TRPV4 channels, as defined by NP_O_ (where N is the number of channels at a site and P_O_ is the open state probability of the channel), was calculated by

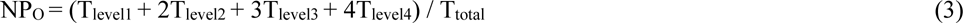

where T is the dwell time at each quantal level detected at TRPV4 sparklet sites and T_total_ is the duration of the recording. NP_O_ was determined using Single Channel Search module of Clampfit and quantal amplitudes derived from all-points histograms ^9^ (ΔF/F_0_ of 0.29 for Fluo-4 –loaded PAs).

Total number of sparklet sites in a field was divided by the number of cells in that field to obtain sparklet sites per cell.

### Immunostaining

Immunostaining was performed on 4^th^-order PAs (∼ 50 μm) pinned *en face* on SYLGARD blocks. PAs were fixed with 4% paraformaldehyde (PFA) at room temperature for 15 minutes and then washed 3 times with phosphate-buffered saline (PBS). The tissue was permeabilized with 0.2% Triton-X for 30 minutes, blocked with 5% normal donkey serum (ab7475, Abcam, Cambridge, MA) or normal goat serum (ab7475, Abcam, Cambridge, MA), depending on the host of the secondary antibody used, for 1 hour at room temperature. PAs were incubated with the primary antibodies (Table 2) overnight at 4°C. Following the overnight incubation, PAs were incubated with secondary antibody 1:500 Alexa Fluor® 568-conjugated donkey anti-rabbit (Life Technologies, Carlsbad, CA, USA) for one hour at room temperature in the dark room. For nuclear staining, PAs were washed with PBS and then incubated with 0.3 mmol/L DAPI (Invitrogen, Carlsbad, CA, USA) for 10 minutes at room temperature. Images were acquired along the z-axis from the surface of the endothelium to the bottom where the EC layer encounters the smooth muscle cell layer with a slice size of 0.1 μm using the Andor microscope described above. The internal elastic lamina (IEL) autofluorescence was evaluated using an excitation of 488 nm with a solid-state laser and collecting the emitted fluorescence with a 525/36 nm band-pass filter. Immunostaining for the protein of interest was evaluated using an excitation of 561 nm and collecting the emitted fluorescence with a 607/36 nm band-pass filter. DAPI immunostaining was evaluated using an excitation of 409 nm and collecting the emitted fluorescence with a 447/69 nm band-pass filter.

**Table 2.**
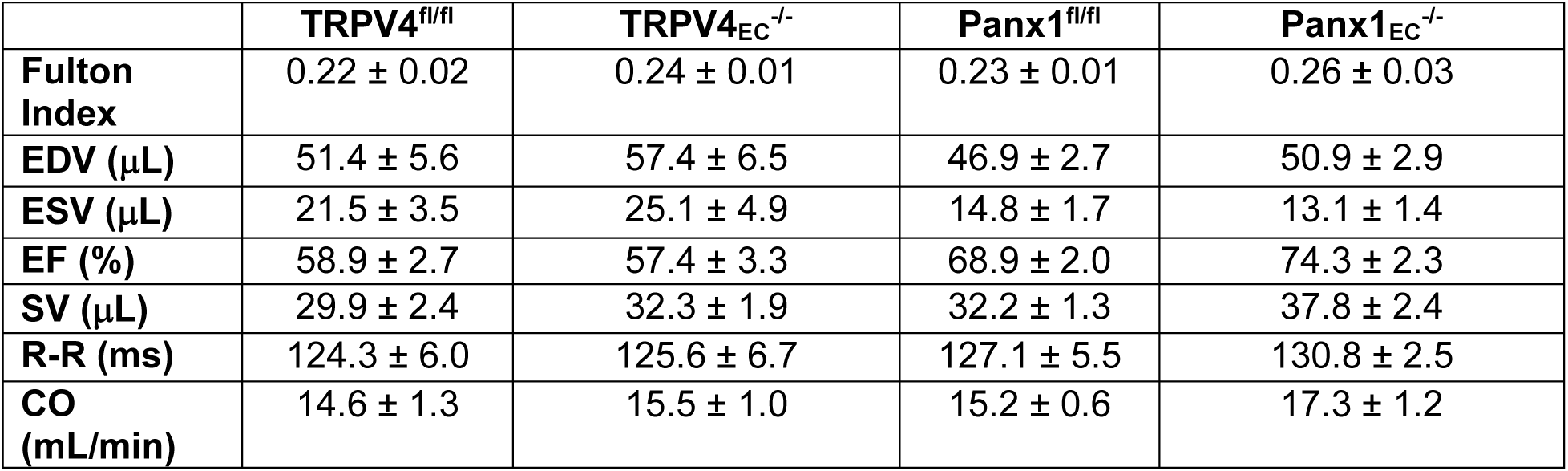
List of antibodies used for immunostaining and PLA on *en face* PAs.

### *In situ* Proximity Ligation Assay (PLA)

Fourth-order (∼ 50 μm) PAs were pinned *en face* on SYLGARD blocks. PAs were fixed with 4% PFA for 15 minutes followed by three washes with PBS. PAs were then permeabilized with 0.2% Triton X for 30 minutes at room temperature followed by blocking with 5% normal donkey serum (Abcam plc, Cambridge, MA, USA) and 300 mmol/L glycine for one hour at room temperature. After three washes with PBS, PAs were incubated with the primary antibodies (Table 2) overnight at 4 °C. The PLA protocol from Duolink PLA Technology kit (Sigma-Aldrich, St. Louis, MO, USA) was followed for the detection of co-localized proteins. Lastly, PAs were incubated with 0.3 mmol/L DAPI nuclear staining (Invitrogen, Carlsbad, CA, USA) for 10 minutes at room temperature in the dark room. PLA images were acquired using the Andor Revolution spinning-disk confocal imaging system along the z-axis at a slice size of 0.1 μm. Images were analyzed by normalizing the number of positive puncta by the number of nuclei in a field of view.

### Plasmid generation and transfection into HEK293 cells

The TRPV4 coding sequence without stop codons was amplified from mouse heart cDNA. The amplified fragment was inserted into a plasmid backbone containing a CMV promoter region for expression and in addition, is suitable for lentiviral production by Gibson assembly. The in-frame FLAG tag was inserted into the 3’-primer used for amplification. Constructs were verified by sequencing the regions that had been inserted into the plasmid backbone. HEK293 cells were seeded (7 x 10^5^ cells per 100 mm dish) in Dulbecco’s Modified Eagle Medium with 10% fetal bovine serum (Thermo Fisher Scientific Inc., Waltham, MA, USA) 1 day prior to transfection. Cells were transfected using the LipofectamineLTX protocol (Thermo Fisher Scientific Inc., Waltham, MA, USA). TRPV4 was co-expressed with PKCα and PKC*β,* obtained from Origene Technologies (Montgomery County, MD).

### Patch clamp in HEK293 cells and freshly isolated ECs

TRPV4 channel current was recorded in HEK293 cells using whole-cell patch configuration 48 hrs after transfection. The intracellular solution consisted of (in mmol/L) 20 CsCl, 100 Cs-aspartate, 1 MgCl_2_, 4 ATP, 0.08 CaCl_2_, 10 BAPTA, 10 HEPES, pH 7.2 (adjusted with CsOH). Currents were measured using a voltage clamp protocol where voltage-ramp pulses (−100 mV to +100 mV) were applied over 200 ms with a holding potential of −50 mV. TRPV4 currents were measured before or 5 minutes after treatment. The extracellular solution consisted of (in mmol/L) 10 HEPES, 134 NaCl, 6 KCl, 2 CaCl_2_, 10 glucose, and 1 MgCl_2_ (adjusted to pH 7.4 with NaOH). Narishige PC-100 puller (Narishige International USA, INC., Amityville, NY, USA) was utilized to pull patch electrodes using borosilicate glass (O.D.: 1.5 mm; I.D.: 1.17 mm; Sutter Instruments, Novato, CA, USA). Patch electrodes were polished using MicroForge MF-830 polisher (Narishige International USA, INC., Amityville, NY, USA). The pipette resistance was (3–5 ΩM). Amphotericin B was dissolved in the intracellular pipette solution to reach a final concentration of 0.3 μmol/L. Data were acquired using HEKA EPC 10 amplifier and PatchMaster v2X90 program (Harvard Bioscience, Holliston, MA, USA), and analyzed using FitMaster v2X73.2 (Harvard Bioscience, Holliston, MA, USA) and MATLAB R2018a (MathWorks, Natick, MA, USA).

Fresh ECs were obtained via enzymatic digestion of 4^th^-order PAs. Briefly, PAs were incubated in the dissociation solution (in mmol/L, 55 NaCl, 80 Na glutamate, 6 KCl, 2 MgCl_2_, 0.1 CaCl_2_, 10 glucose, 10 HEPES, pH 7.3) containing Worthington neutral protease (0.5 mg/mL) for 30 minutes at 37°C. The extracellular solution consisted of (in mmol/L) 10 HEPES, 134 NaCl, 6 KCl, 2 CaCl_2_, 10 glucose, and 1 MgCl_2_ (adjusted to pH 7.4 with NaOH). The intracellular pipette solution for perforated-patch configuration consisted of (in mmol/L) 10 HEPES, 30 KCl, 10 NaCl, 110 K-aspartate, and 1 MgCl_2_ (adjusted to pH 7.2 with NaOH). Cells were kept at room temperature in a bathing solution consisting of (in mmol/L) 10 HEPES, 134 NaCl, 6 KCl, 2 CaCl_2_, 10 glucose, and 1 MgCl_2_ (adjusted to pH 7.4 with NaOH). TRPV4 channel current was recorded from freshly isolated ECs as described previously ^25, 61^. Briefly, GSK101-induced outward currents through TRPV4 channels were assessed in response to a 200-ms voltage step from −45 mV to +100 mV in the presence of ruthenium red in order to prevent Ca^2+^ and activation of IK/SK channels at negative voltages. Outward currents were obtained by averaging the currents through the voltage step. GSK219-sensitive currents were obtained by subtracting the currents in the presence of GSK219 from the currents in the presence of GSK101.

### Statistical analysis

Results are presented as mean ± SEM. The n=1 was defined as one artery in the imaging experiments (Ca^2+^ imaging, PLA), one cell for patch clamp experiments, one mouse for RVSP measurements, one artery for pressure myography experiments, one mouse for functional MRI, one mouse for ATP measurements, and one mouse for qPCR experiments. The data were obtained from at least five mice in experiments performed in at least two independent batches. The individual data points are shown each dataset. For *in vivo* experiments, an independent team member performed random assignment of animals to groups and did not have knowledge of treatment assignment groups. All the *in vivo* experiments were blinded; information about the groups or treatments was withheld from the experimenter or from the team member who analyzed the data.

All data are shown in graphical form using CorelDraw Graphics Suite X7 (Ottawa, ON, Canada) and statistically analyzed using GraphPad Prism 8.3.0 (Sand Diego, CA). A power analysis to determine group sizes and study power (>0.8) was performed using GLIMMPSE software (α = 0.05; >20% change). Using this method, we estimated at least 5 cells per group for patch clamp experiments, 5 arteries per group for imaging and pressure myography experiments, and 4 mice per group for RVSP measurements and MRI. A Shapiro-Wilk test was performed to determine normality. The data in this article were normally distributed; therefore, parametric statistics were performed. Data were analyzed using two-tailed, paired or independent t-test (for comparison of data collected from two different treatments), one-way ANOVA or two-way ANOVA (to investigate statistical differences among more than two different treatments). Tukey correction was performed for multiple comparisons with one-way ANOVA, and Bonferroni correction was performed for multiple comparisons with two-way ANOVA. Statistical significance was determined as a P value less than 0.05.

## Acknowledgements

The mouse strain cdh5-optoα1AR was developed by CHROMus^TM^ which is supported by the National Heart Lung Blood Institute of the National Institute of Health under award number R24HL120847.

